# Read depth correction for somatic mutations

**DOI:** 10.1101/2022.02.16.480761

**Authors:** Jordan Anaya, Alexander S. Baras

**Affiliations:** Department of Pathology, Johns Hopkins University School of Medicine, Baltimore, MD, USA; The Sidney Kimmel Comprehensive Cancer Center, Johns Hopkins University School of Medicine, Baltimore, MD, USA; Bloomberg Kimmel Institute for Cancer Immunotherapy, Sidney Kimmel Comprehensive Cancer Center, Johns Hopkins University School of Medicine, Baltimore, MD, USA

**Author notes:** Corresponding author: Alexander S. Baras.

## Abstract

The ability to accurately detect mutations is a function of read depth and variant allele frequency (VAF). While the read depth distribution of a sample is observable, the true VAF distribution of all mutations in a sample is uncertain when there is low coverage depth. We propose to estimate the VAF distributions that would be observed with high-depth sequencing for samples with low sequencing depth by grouping samples with similar clonality and purity and using the VAF distributions observed with the high-depth mutations that are available. With these estimated high-depth VAF distributions we then calculate what the expected VAF distributions would be at a given depth and compare against the observed VAF distributions at that depth. Using this procedure we estimate that The Cancer Genome Atlas (TCGA) MC3 dataset only reports on average 83% of the mutations in a sample which would have been detected with high-depth sequencing. These results have important implications for comparing tumor mutational burden (TMB) estimates when samples are sequenced at different depths and for modeling high-depth, gene panel-based sequencing from the TCGA MC3 dataset.

## INTRODUCTION

While it is common to normalize some omics data by depth of sequencing^1^, to our knowledge the use of read depth with somatic mutations has been limited to quality control metrics and in constructing a metric for the prevalence of a mutation in a given sample (i.e. VAF). Models that use an aggregation of mutation counts, for example a model that predicts tumor type^2^, should apply some correction for variability in read depth, and ideally also take the uncertainty of the correction into account. Given the recent FDA approval of immunotherapy for patients with TMB ≥ 10 as defined by the FoundationOneCDx assay^3,4^, and recent proposals to calibrate panel-based TMB to TCGA exome data^5,6^, there is an urgent need to characterize how differences in coverage depth may affect estimates of TMB.

It is generally understood that at read depths typically seen in whole exome or whole genome sequencing, approximately 50-100X, mutation callers will have a considerable false negative rate^7,8^. While it may seem possible to use the performance characteristics of different callers to estimate how many mutations will be missed in a sample, the performance depends on the VAF and depth of sequencing and the true VAF distribution is not known since mutations are potentially missing. In addition, the mutations in a given sample may not be enough to produce a smooth distribution, VAFs which are observed at low depths will be noisy measures, and at low depths there are mathematical and practical limits on the minimum VAFs that will be seen.

Although we cannot infer the exact VAF distribution of a sample, we can estimate it by grouping samples together that we think should have similar VAF distributions, and then determine the true underlying VAF distributions by only using mutations sequenced at a depth seen in panel sequencing (Figure 1A). It is known that the VAF distribution depends on the purity, clonality, and ploidy of a sample. In fact, clonality can be determined if VAF, purity, and copy number are known^9^. Consistent with this, we observed markedly different VAF distributions when we grouped samples by their purity and clonality (Figure 1B). It’s possible that there’s a third axis (copy number alterations) that should be taken into account, but confidence in the distributions requires a certain amount of data which may no longer be present with additional splitting.

**Figure 1.**
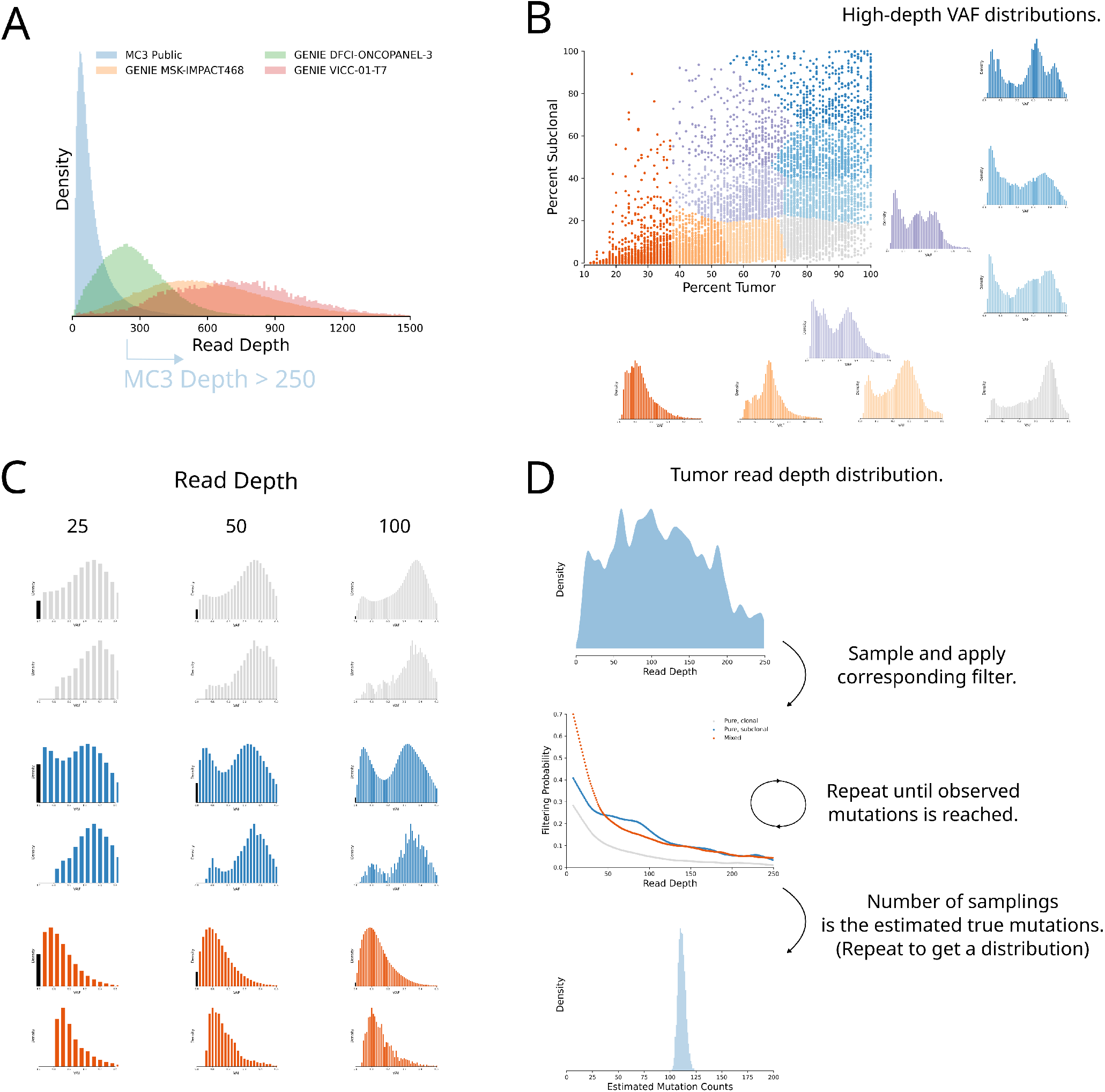
Read depth correction procedure. (A) The depth at observed mutations for several different assays. (B) Scatter plot of TCGA MC3 samples with the colors corresponding to the assigned k-means clusters derived from purity and clonality values. For each cluster of samples the VAF distribution of mutations at a depth greater than 250 is shown. (C) Theoretical VAF distributions for different clusters and different read depths (top) and the corresponding observed distributions at those depths (bottom). The zero bin is shown in black. The difference between the distributions is the assumed probability that a mutation wasn’t called at that depth. (D) To estimate the true number of mutations in a sample the read depth distribution is sampled and then potentially filtered according to the corresponding probability curve. This process is repeated until the number of samplings which passed the filter equals the number of observed mutations. Repeating this process generates a distribution of possible true mutation counts for a sample (observed mutations was 100).

If we take these high-depth VAF distributions from grouped samples to be the underlying true distribution for each individual sample in a group, then it is straightforward to generate the exact VAF distribution we would expect at a given read depth for a sample (Figure 1C). This distribution will include 0, and those bins are highlighted in black. We can then look at what VAF distribution was observed at that read depth for the samples in that group (Figure 1C). As expected, all observed distributions are missing the black bin given it is impossible to call a mutation with 0 variant reads, but there are other noticeable differences between the distributions, particularly at lower VAFs. These differences can be either due to missing data, or alternatively, additional data. This additional data would have to be falsely called mutations, and since they would have to be at higher VAFs would presumably be germline mutations which were accidentally called as somatic. However, because we had strong a priori reasons to expect missing data at lower VAFs and have confidence in the MC3 working group’s filtering of germline mutations, we will assume the differences between distributions are the result of mutations which were missed due to a complex relationship between the mutation calling pipeline and quality of the sequencing data.

By assuming data is missing we can then add mass to the observed distribution with the goal of making it appear similar to the expected distribution, and the resulting ratio of the amount of data added to the total amount of data in the final distribution represents the chance of missing a mutation at that read depth. This process will generate a probability of missing mutations at every read depth not considered to be high (above 250 in this case) for each group (Figure 1D). As expected the samples which are pure and clonal had lower filtering probabilities than either pure subclonal samples or mixed samples.

To get a read depth distribution for each sample we randomly sampled the BAMs at 1000 locations in the exome, and used the resulting counts to generate kernel densities, one of which is shown in Figure 1D. To estimate how many mutations a sample would have if it were sequenced at higher depth we randomly sample from this distribution and then use the corresponding probability of missing a mutation at the sampled depth to determine if a mutation was called. This process is continued until the number of called mutations matches the number of observed called mutations. In Figure 1D we show the results of this process repeated 10,000 times for a sample with an observed mutation count of 100. The mean and shape of this distribution will be dependent on a sample’s read depth distribution and its corresponding filtering probability curve.

## VALIDATION

We are in the early stages of assessing the accuracy of our estimates, but we were interested in determining whether our current estimates added any value. In addition to undergoing exome sequencing, a certain number of TCGA samples also underwent genome sequencing as part of the Pan-Cancer Analysis of Whole Genomes (PCAWG) Consortium. Recently the mutation calls between these two consortia were compared^10^. The comparison identified mutations which were only called by the MC3 working group, mutations which were only called by the PCAWG working group, and called by both (Figure 2A). Of course, there are in theory mutations which were called by neither group. If an MC3 sequenced sample failed to call a large fraction of mutations then there should be a higher chance of the PCAWG group detecting new mutations. Conversely, if the MC3 group called every possible mutation then the PCAWG sequencing should not detect any new mutations. As a result, we would expect there to be a correlation between the ratio of MC3 called mutations to all called mutations and our estimates of what percent of mutations were observed (Figure 2B). We did observe a strong positive correlation (Pearson’s r = .45, p = 1.03e-32), which was much stronger than the correlation with purity values (Pearson’s r = .23, p = 3.70e-9). It is unclear what correlation would be achieved with perfect predictions due to the uncertainty associated with the target metric.

**Figure 2.**
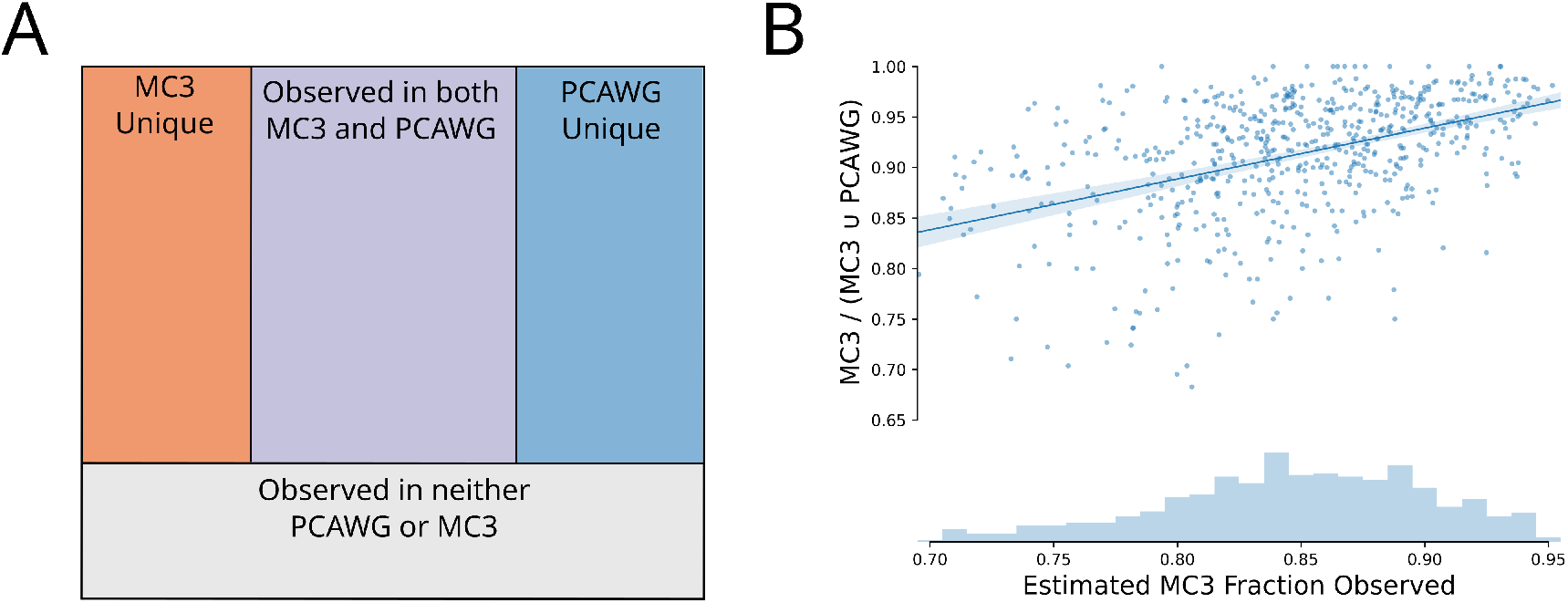
Validation with PCAWG re-sequencing. (A) Illustration of the possible outcomes of re-sequencing TCGA samples by the two working groups. (B) Scatter plot of our fraction observed estimates to the ratio of number of mutations called by the MC3 working group and all called mutations.

## DISCUSSION

It was recently proposed that the TMB of low purity samples needs to be adjusted^11^. While we agree that low purity samples typically need a larger correction factor than higher purity samples, we suggest that samples with coverage depth lower than 150X would strongly benefit from adjustment, regardless of purity. This has implications for TMB calibration given that conventional exome sequencing (which is currently viewed as the gold standard measure for TMB) has a wide distribution of coverage depth both within and between samples. When this variability is not accounted for it adds potential noise to the target exomic TMB. Although we currently don’t have a pipeline set up to assess the accuracy of our estimates, when panel data was downsampled to 100X only 83.5% of mutations were re-called, which is in line with our estimates for the MC3 data^12^.

The true underlying VAF distribution not only depends on the purity and clonality of the samples, but also on the mutation calling pipeline. For example, the controlled MC3 MAF reports all variants which were called by any caller (the public MAF requires 2 callers), which results in more mutations at lower VAFs. These lower VAFs bias all of the expected distributions to lower VAFs, resulting in a concurrent increase in filtering probabilities at every read depth. As a result, when using the MC3 VAF distributions for your data it is essential that your processing pipeline is expected to generate similar VAF distributions. If your pipeline used a single mutation caller it may be possible to limit the MC3 data to that mutation caller and then use the resulting distributions. Or if your pipeline includes filters such as excluding all mutations below a certain VAF it should be possible to apply those filters to the true, expected, and observed distributions and generate new probability curves.

When calibrating TMB it may seem unnecessary to normalize the MC3 data (essentially upscale the data) as long as every panel is calibrated to the same target. However, not every dataset will be panel data. For example, clinical trials that use exomic sequencing will view their values as the true values, but if they have high-depth data their values will be higher than values calibrated to the MC3 dataset. Although we haven’t assessed the accuracy of our estimates and our algorithm is evolving, we would recommend that researchers using somatic mutation counts consider the effect read depth may have on their results and ways to correct for it.

## METHODS

### Availability of Data and Materials

All data used in this publication is publicly available, albeit the clonality MAF and BAMs are controlled access. The MC3 MAF is from [13]. The GENIE 10.1 mutations and panel information are available at Synapse, https://www.synapse.org/#!Synapse:syn25895958^14^. Purity information was downloaded from https://gdc.cancer.gov/about-data/publications/pancanatlas/. A MAF with clonality predictions was downloaded from https://gdc.cancer.gov/about-data/publications/pancan-aneuploidy/. The GFF used was downloaded from ftp://ftp.ensembl.org/pub/grch37/current/gff3/homo_sapiens/Homo_sapiens.GRCh37.87.gff3.gz. The BED for the Broad’s Agilent kit was downloaded from https://bitbucket.org/cghub/cghub-capture-kit-info/src/master/BI/vendor/Agilent/whole_exome_agilent_1.1_refseq_plus_3_boosters.targetIntervals.bed. Coverage WIGs were downloaded from https://www.synapse.org/#!Synapse:syn21785741. All code for the results in this manuscript is available at GitHub: https://github.com/OmnesRes/depth_norm, and has been archived at Zenodo: https://doi.org/10.5281/zenodo.6081107. Code is written in Python 3.

### TCGA MC3 MAF Processing

Variants in the MC3 public MAF which had a “FILTER” value of either “PASS”, “wga”, or “native wga mix” were retained. Coverage WIGs were converted to BEDs with wig2bed. Variants were dropped if they didn’t fall in the corresponding tumor/normal coverage coordinates.

### Read Depth Distribution

To get an unbiased distribution of read depths for each sample we randomly selected 1000 locations from the Broad’s Agilent kit limited to the CDS via PyRanges^15^. We then lifted over the coordinates from GRCh37 to GRCh38 and constructed queries for the genomic data portal’s BAM slicing API. We then used pysam to count the number of nonrefskips at each of the 1000 locations. Because it’s impossible to know if a location with no reads is missing reads because of a lack of depth of sequencing or a lack of coverage of the exome kit, only counts with at least minimum depth observed in the MC3 calls were used. The kernel density was also only calculated for counts under what we consider to be high depth (250). Above this we cutoff we assume mutations will always be detected.

### Assumed VAF Distributions

Purity and subclonality values were assigned to each sample. Subclonality was defined as the percent of mutations in a sample which were predicted to be subclonal. Because it is difficult to determine clonality below a certain purity, all samples with a purity of 37% or less were considered a single group and excluded from k-means clustering. The VAFs of high-depth mutations in each group were rounded to 2 decimals (essentially creating 101 bins). To generate an expected VAF distribution at a given read depth a binomial distribution was calculated for each of the 101 VAFs and then the final distribution was weighted by the proportion of each VAF in the high-depth distribution.

### Filtering Probabilities

Observed distributions were constructed by looking at the actual mutations for each group at a given read depth. For some depths there is a limited amount of data and in these cases mutations of neighboring depths are also used. To determine how much data needed to be added to the observed distributions data was added iteratively with the constraint that data was only added to bins at lower VAFs. An observed distribution has an expected noise associated with it, and this uncertainty was taken into account when adding data. Probabilities were smoothed with lowess.

### PCAWG Validation

The variant-level filter was set to “PASS” for MAFit (https://mbailey.shinyapps.io/MAFit/). Samples which had only had 5 or fewer shared mutations were discarded. For the purposes of estimating what fraction of a sample’s mutations were detected each sample was assumed to have 100 observed mutations and the average of 100 estimates was treated as the point estimate. These estimates for all TCGA samples were also calculated and are made available as supplemental data.

## ACKNOWLEDGMENTS

The results here are in whole or part based upon data generated by the TCGA Research Network.

